# Conditioned immunological memory: implication for cognition of immune cells

**DOI:** 10.1101/2024.07.14.603476

**Authors:** Zhu Zhu, Wei Qin, Qianlin Li, Hantao Zhang, Zixuan Lin, Lin Chen, Yunan Zhao

**Affiliations:** Department of Pathology and Pathophysiology, School of Medicine, Nanjing University of Chinese Medicine, Nanjing 210023, China; Department of Physiology, School of Medicine, Nanjing University of Chinese Medicine, Nanjing 210023, China

**Keywords:** associative learning, immunological memory, conditioned stimulus, cognition, hapten

## Abstract

Associative learning refers to the process by which animals or human beings build the new relationship between two stimuli or events, which has been found throughout the animal kingdom. However, no studies have reported the associative learning in cells in the body to date. In the study, we focused on immune cells, as immunological memory is considered as one of the cardinal features of the immune system. As for a classical conditioning procedure, we designed the analogues of ginsenoside Rh1 (e.g., protopanaxadiol and panaxadiol) as the conditioned stimulus (CS) and the mcKLH-Rh1 conjugate as the unconditioned stimulus (US) and determined the CS-US association by detecting the dynamics of anti-Rh1 antibody production in serum. To test the hypothesis that immune cells also have the same associative learning ability as animals or human beings, mice were divided into control and CS group. Based on above CS and US, two groups of mice were subjected to conditioned learning, conditional memory recall, and deconditioning. When we used protopanaxadiol as the CS, the antibody response to Rh1 in the mice of CS group was slower and weaker than the control mice during conditional memory recall. Interestingly, the difference between two groups in antibody response to the immunogen disappeared during deconditioning. When we used panaxadiol as the CS, the antibody response to Rh1 in the mice of two groups was similar with that in studies regarding protopanaxadiol as the CS throughout three experimental phases. These results suggested that immune cells could acquire new association between a hapten analogue and a hapten-protein conjugate after undergoing a classical conditioning. Moreover, the conditioned immunological memory enables immune cells to make a conditioned response of humoral immunosuppression to prevent US induced immune damage. Our findings open a new frontier to explore cognitive functions of cells in the body and contribute to a novel understanding of cellular bioactivities.

## 1. Introduction

Learning can be generally divided into two categories: non-associative learning[1], in which animals learn about the occurrence or properties of a single event or stimulus, and associative learning[2], in which animals acquire the new association between two temporally discrete events or stimuli. During a classical conditioning procedure of associative learning, one conditioned stimulus (CS) is repeatedly followed by another unconditioned stimulus (US). After successfully building the association, animals make a conditioned response to the CS that deal with the upcoming US, and it is generally thought to have learned that the CS predicts the arrival of the US.

For example, conditioned fear memory represents the formation of an association between a neutral conditioned stimulus (e.g., a tone, context, or odor) and an unconditioned aversive or punitive stimulus (e.g., a foot-shock or loud noise). Following conditioning, the presentation of the CS alone evokes a defensive response (e.g., freezing of movement or escape behavior)[3]. The defensive response is stable and results from retrieval of the stored fear memory. Consequently, the acquired CS-US association enables animals to predict aversive or punitive events, which is an adaptive process to prevent harm and help animals survive in the constantly changing environment[4]. Nowadays, the properties of classical conditioning have been studied in a wide range of species and have been found to similar throughout the animal kingdom. Moreover, associative learning relates to animal cognition[5], and predicts intelligence above and beyond working memory and processing speed in human beings[6], which plays a variety of roles in the study of animal or human cognition.

As a traditional medicine, Chinese medicine has a long history and still plays an important role in health care in China. Based on over 2000 years of practice, Chinese medicine has a lot of unique principles (e.g., Yin and Yang, Qi, five basic elements, and six pathogenic factors)[7, 8], which was mainly derived from a sensual, figurative, and intuitive understanding of the natural world. Ancient Chinese scholars then applied such knowledge to understand all kinds of complex things by analogy or symbolism. In ancient times, Chinese medicine practitioners could not explore the mysteries of human life involving biology, physiology and pathology without the help of advanced technologies as today. However, they believe that the internal active principles of human body are like those of the natural world or human society (e.g., mutually reinforcing and mutually restricting among the internal organs)[9], and then they apply such principles summarized from the nature world or human society to understand, prevent, and cure diseases (e.g., Monarch, Minister, Assistant, and Guide in the composition of prescription and certain function imbalance of organs while getting sick)[10]. Inspired by the mindset of ‘abstraction and analogy’ in Chinse medicine, we proposed a hypothesis that cells in the body also have the same associative learning ability as animals or humans, since associative learning has been found throughout the animal kingdom.

To test this hypothesis, in the study, we used natural compounds as the CS and an immunogenic conjugate (hapten-protein) as the US to determine whether it was possible for immune cells to successfully build the CS-US association, confirming that cells in the body also have cognitive abilities.

## 2. Materials and Methods

### 2.1 Chemicals

The ginsenosides Rh1, Rb1, Rd, protopanaxadiol (PPD), and panaxadiol (PD) analytical standards (purity ≥ 98%) were purchased from Shanghai yuanye Bio-Technology Co., Ltd (Shanghai, China) (Figure 1). Imject^TM^ Mariculture keyhole limpet hemocyanin (mcKLH), Imject^TM^ Freund’s complete adjuvant (FCA), Imject^TM^ Freund’s incomplete adjuvant (FIA), and Horseradish peroxidase (HPR)-conjugated goat anti-mouse IgG (H+L) secondary antibody were supplied by Thermo Fisher Scientific Inc. (Loughborough, UK). Deionized water was prepared using a Milli-Q water purification system (Millipore, Billerica, MA, USA). Bovine serum albumin (BSA), 3,3’,5,5’-Tetramethylbenzidine (TMB) two-component substrate solution, and ELISA stop solution were purchased from Solarbio Science & Technology Co., Ltd (Beijing, China). All other analytical grade solvents were obtained from Nanjing Chemical Reagents Co. (Nanjing, China).

**Figure 1.**
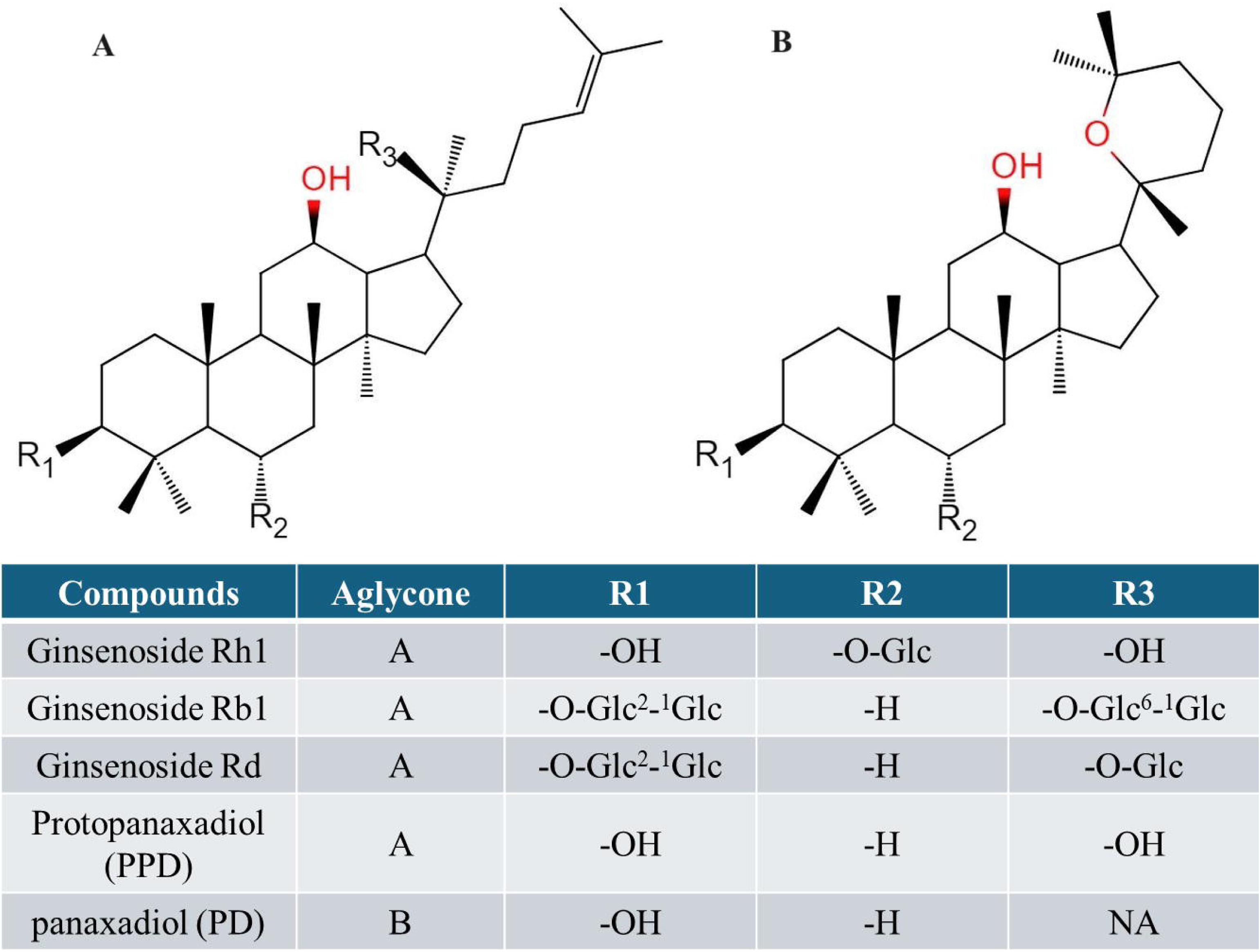
Structures of ginsenosides and aglycones. Rh1, Rb1, and Rd differ at 3 side chains attached to the common aglycone (protopanaxadiol, PPD). Panaxadiol (PD) has the same steroid ring as PPD, but they differ at the side chain. The meaning of abbreviation (Glc) is glucopyranoside.

### 2.2 Animals

Male C57BL/6N mice weighing 18-22 g each were obtained from SLAC Laboratory Animal Co., Ltd. (Shanghai, China). These mice were maintained under a standard 12-h light/dark cycle (lights on between 7:00 and 19:00) with ad libitum access to food and water at a constant temperature (22 ± 2 °C). They were housed in groups of five per cage and adapted to daily handling after delivery. All animal procedures were approved by the IACUC (Institutional Animal Care and Use Committee) of Nanjing University of Chinese Medicine (License Number: 202012A042) and carried out in accordance with the Guidelines of Accommodation and Care for Animals. We used the minimum number of mice required to obtain consistent data.

### 2.3 Preparation of immunogenic (hapten-protein) conjugates

Rh1-protein conjugates were synthesized as previously described[11] (Figure 2). Briefly, a solution of Rh1 (5 mg) in ethanol (0.7 mL) was added dropwise to a solution of NaIO_4_ (4 mg, dissolved with 0.5 mL of deionized water) over 1 h under stirring at room temperature. Then, the mixture was added to a solution of BSA (10 mg) in 100 mM carbonate buffer (pH 9.2, 1 mL) or a dsolution of mcKLH (10 mg) in phosphate-buffered saline (PBS, 1 mL), which was stirred at room temperature for 2 h. Subsequently, a fresh solution of NaBH_4_ (3 mg) in deionized water (0.5 mL) was added dropwise to the mixture, which was then incubated and shaken at room temperature for 3 h. Finally, the reaction mixture was dialyzed against deionized water at 4 °C for 2 days using Slide-A-Lyzer^TM^ Dialysis Cassette (10K MWCO, 3 mL, Thermo Scientific) and stored at −70 °C.

**Figure 2.**
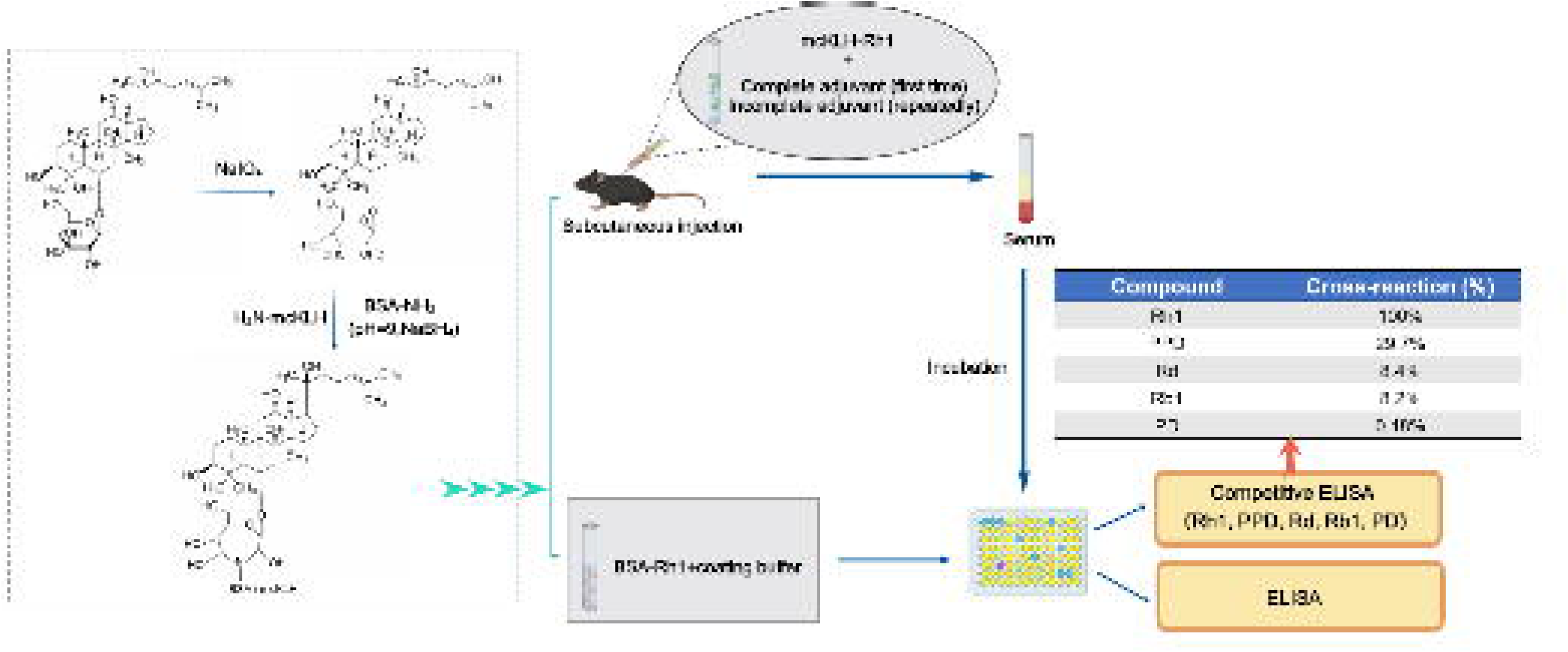
Flowchart showing the preparation of protein-Rh1 conjugates, immunization, and immunoassays of ELISA and competitive ELISA.

### 2.4 Immunization using mcKLH-Rh1 conjugate

Male C57BL/6N mice were used for immunization with mcKLH-Rh1 conjugate. For the first injection, mcKLH-Rh1 conjugate was emulsified with FCA in the same ratio. This emulsion (40 µg protein) was subcutaneously administered into a mouse. The same dose of the immunogen mixed in the same ratio with FIA was used as a booster once every 2 weeks for 1-2 months. The immune serum was collected after the final immunization for direct ELISA and competitive indirect ELISA.

### 2.5 Enzyme-linked immunosorbent assay (ELISA) using BSA-Rh1 conjugate

A 96-well microplate was coated with a solution of BSA-Rh1 conjugate (100 µL, 1 mg/mL) in 50 mM carbonate buffer (pH 9.6) overnight at 4 °C. After washing the plate 4 times with PBS containing 0.05% Tween 20 (PBST), 50 µL of PBS, diluted negative serum (1:10), and diluted immune serum (1:10) were added into each well and incubated for 1 h at 37 °C. After washing as above, 50 µL of diluted HPR-conjugated goat anti-mouse secondary antibody (1:10,000) was added to each well and incubated for 30 min at 37 °C. After washing, 100 µL of TMB substrate solution was added to each well. Following incubation for 10 min at 37 °C, the reaction was stopped by adding 100 µL of ELISA stop solution. The absorbance was measured at 450 nm using an ELISA reader (SpectraMax M5, Molecular Devices, USA). The reactivity of immune serum to BSA-Rh1 conjugate was expressed as P/N value [P/N = A_450_(immune serum-blank)/A_450_(negative serum-blank)]. The mean value of the triplicate readings for each sample was calculated.

### 2.6 Competitive ELISA for cross-reactivity determination of immune serum against Rh1

A 96-well microplate was coated with 100 µL of BSA-Rh1 conjugate (1 mg/mL) in 50 mM carbonate buffer (pH 9.6) overnight at 4 °C. After washing 4 times with PBST, 50 µL of mixture was added into each well and incubated for 1 h at 37 °C. The mixture was prepared by mixing 10 times-diluted immune serum with a solution of compound (Rh1, Rd, Rb1, PPD or PD, 2 µM in PBS containing 0.01% dimethyl sulfoxide) or the vehicle in the same ratio. After washing as above, 50 µL of diluted HPR-conjugated goat anti-mouse secondary antibody was added to each well and incubated for 30 min at 37 °C. After washing, 100 µL of TMB substrate solution was added to each well. Following incubation for 10 min at 37 °C, the reaction was stopped by adding 100 µL of ELISA stop solution. Subsequently, the absorbance was measured at 450 nm using an ELISA reader. The cross-reactivity of immune serum against Rh1 was expressed as cross-reaction ratio (%) [A_450_(vehicle-tested compound) × 100/A_450_(vehicle-Rh1)]. The data were obtained with triplicates.

### 2.7 Conditioned immunological memory

As depicted in Figure 3, mice were divided into two groups (n = 5/group): control group and conditioned stimulus group. The experimental procedures consisted of three phases (conditioned learning, conditional memory recall, and deconditioning) and lasted for approximately one year.

**Figure 3.**
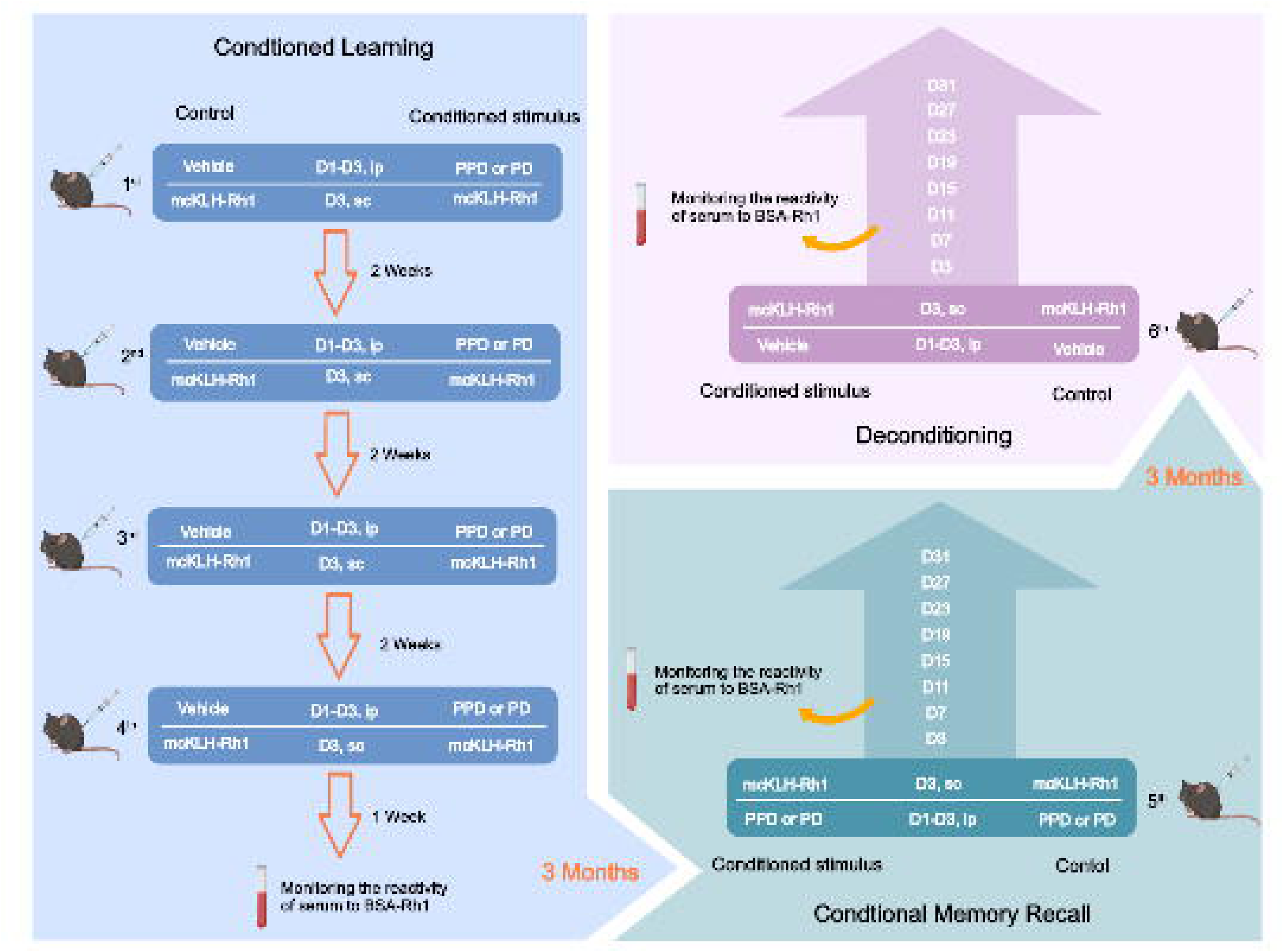
General experimental procedure for conditioned immunological memory. Mice were divided into two groups: control group and conditioned stimulus group. These two groups of mice were subjected to three experimental phases: conditioned learning, conditional memory recall, and deconditioning. The whole procedure lasted for approximately one year.

#### 2.7.1 Conditioned learning

The first experimental phase lasted for approximately two months. During which, the mice of two groups were submitted to immunization with mcKLH-Rh1 conjugate (40 µg per mouse) once every 2 weeks for a total of 4 times. As for each immunization, the mice of the conditioned stimulus group were injected intraperitoneally with PPD or PD (20 mg/kg, suspended in PBS containing 0.1% dimethyl sulfoxide) once a day at random times during light phase for three days (day 1-day 3). The dosage of PPD or PD was based on previous pharmacokinetic reports that it could produce a serum peak concentration of 1-10 μM in rodent animals after intraperitoneal administration[12–14]. The control mice were injected intraperitoneally only with the vehicle. Half an hour after the intraperitoneal injection on the third day (day 3), the mice of two groups were subcutaneously administered mcKLH-Rh1 conjugate emulsified with FCA or FIA in the same ratio. One week after the final immunization, blood samples (50 µL) were collected from the lateral saphenous vein of the mice. Serum was separated by centrifugation at 2500 rpm for 5 min and stored at −70 °C until ELISA.

#### 2.7.2 Conditional memory recall

3∼4 months after the first experimental phase, conditional memory recall was carried out, which lasted for approximately one month. During the second experimental phase, the mice of two groups were injected intraperitoneally with PPD or PD (20 mg/kg, suspended in PBS containing 0.1% dimethyl sulfoxide) once a day at random times during light phase for three days (day 1-day 3). Half an hour after the intraperitoneal injection on the third day (day 3), the mice of two groups were subcutaneously administered mcKLH-Rh1 conjugate (40 µg per mouse) emulsified with FIA in the same ratio. After the immunization, blood samples (50 µL) were collected from the lateral saphenous vein of the mice every four days for a total of 8 times (day 3, day 7, day 11, day 15, day 19, day 23, day 27, and day 31). Serum was separated by centrifugation at 2500 rpm for 5 min and stored at −70 °C until ELISA.

#### 2.7.3 Deconditioning

3∼4 months after the second experimental phase, deconditioning was performed, which lasted for approximately one month. During the third experimental phase, the mice of two groups were injected intraperitoneally with the vehicle once a day at random times during light phase for three days (day 1-day 3). Half an hour after the intraperitoneal injection on the third day (day 3), the mice of two groups were subcutaneously administered mcKLH-Rh1 conjugate (40 µg per mouse) emulsified with FIA in the same ratio. After the immunization, blood samples (50 µL) were collected from the lateral saphenous vein of the mice every four days for a total of 8 times (day 3, day 7, day 11, day 15, day 19, day 23, day 27, and day 31). Serum was separated by centrifugation at 2500 rpm for 5 min and stored at −70 °C until ELISA.

### 2.8 Statistical analysis

Data were expressed as mean ± standard error of mean (S.E.M.) and analyzed using GraphPad Prism 10. As for comparing multiple groups, statistical significance was determined by repeated measures two-way analysis of variance (ANOVA) followed by Turkey’s post hoc test. As for comparing two groups, a two-tailed Student’s t-test was used. The significance level was set at *P*≤0.05 for all statistical comparisons.

## 3. Results

### 3.1 Associative learning of immune cells between PPD and mcKLH-Rh1 conjugate

As depicted in Figure 4A, first, there was no significant difference for the reactivity of immune serum against Rh1 between two groups at the end of conditioned learning. Second, during conditional memory recall, two-way ANOVA showed significant effects of time (*F*_7, 56_ = 14.740; *P* < 0.01) and conditioned stimulus (*F*_1, 8_ = 13.500; *P* < 0.01) on the reactivity of immune serum against Rh1 with a significant interaction between two factors (*F*_7, 56_ = 6.549; *P* < 0.001). Multiple comparisons test showed that the reactivity of immune serum against Rh1 in the mice of conditioned stimulus group significantly decreased at day 7 (*P* < 0.001), day 11 (*P* < 0.01), day 15 (*P* < 0.01), day 19 (*P* < 0.05), day 23 (*P* < 0.05), day 27 (*P* < 0.05), and day 31 (*P* < 0.05), but not at day 3. Meanwhile, the area under the curve (AUC) of serum antibody response to Rh1 in the mice of conditioned stimulus group also significantly decreased in contrast to the control group (Figure 4B). Third, after deconditioning, there was no significant difference for the AUC of serum antibody response to Rh1 between two groups (Figure 4C). There was also no significant effect of conditioned stimulus (*F*_1, 8_ = 1.573; *P* = 0.245) and the interaction between time and conditioned stimulus (*F*_7, 56_ = 0.406; *P* = 0.895) on the reactivity of immune serum against Rh1. The effect of time, however, was statistically significant (*F*_7, 56_ = 10.660; *P* < 0.01).

**Figure 4.**
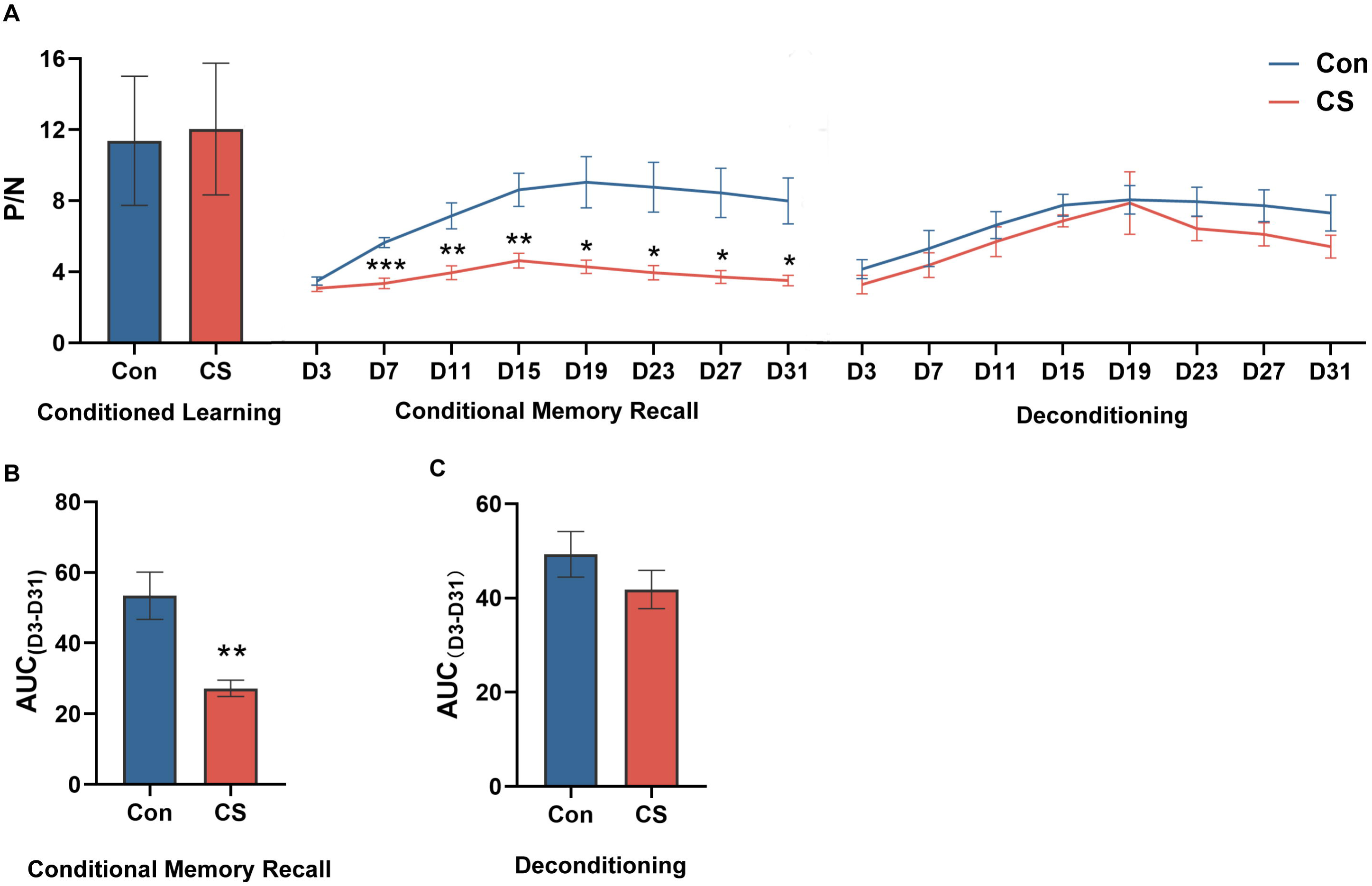
Associative learning of immune cells between PPD and mcKLH-Rh1 conjugate. (A) The reactivity of immune serum to BSA-Rh1 conjugate (P/N) in the process of conditioned learning, conditional memory recall, and deconditioning. (B) Areas under curve (AUC) of P/N during conditional memory recall. (C) Areas under curve (AUC) of P/N during deconditioning. *p < 0.05, **p < 0.01 and ***p < 0.001 vs control group; mean ± SEM (n = 5).

### 3.2 Associative learning of immune cells between PD and mcKLH-Rh1 conjugate

As depicted in Figure 5A, first, there was no significant difference for the reactivity of immune serum against Rh1 between two groups at the end of conditioned learning. Second, during conditional memory recall, two-way ANOVA showed a significant effect of time (*F*_7, 56_ = 6.011; *P* < 0.05) and an almost significant effect of conditioned stimulus (*F*_1, 8_ = 5.003; *P* = 0.056) on the reactivity of immune serum against Rh1 with no significant interaction between two factors (*F*_7, 56_ = 1.321; *P* = 0.258). Multiple comparisons test showed that the reactivity of immune serum against Rh1 in the mice of conditioned stimulus group trended to decrease at day 7 (*P* = 0.085), day 15 (*P* = 0.087), and day 23 (*P* = 0.054). Whereas the AUC of serum antibody response to Rh1 in the mice of conditioned stimulus group significantly decreased in contrast to the control group (Figure 5B). Third, after deconditioning, there was no significant difference for the AUC of serum antibody response to Rh1 between two groups (Figure 5C). Two-way ANOVA also showed that there was no significant effect of conditioned stimulus (*F*_1, 8_ = 0.950; *P* = 0.358) on the reactivity of immune serum against Rh1 with no significant interaction between time and conditioned stimulus (*F*_7, 56_ = 1.243; *P* = 0.295). There was, however, a significant effect of time on the reactivity of immune serum against Rh1 (*F*_7, 56_ = 12.000; *P* < 0.001).

**Figure 5.**
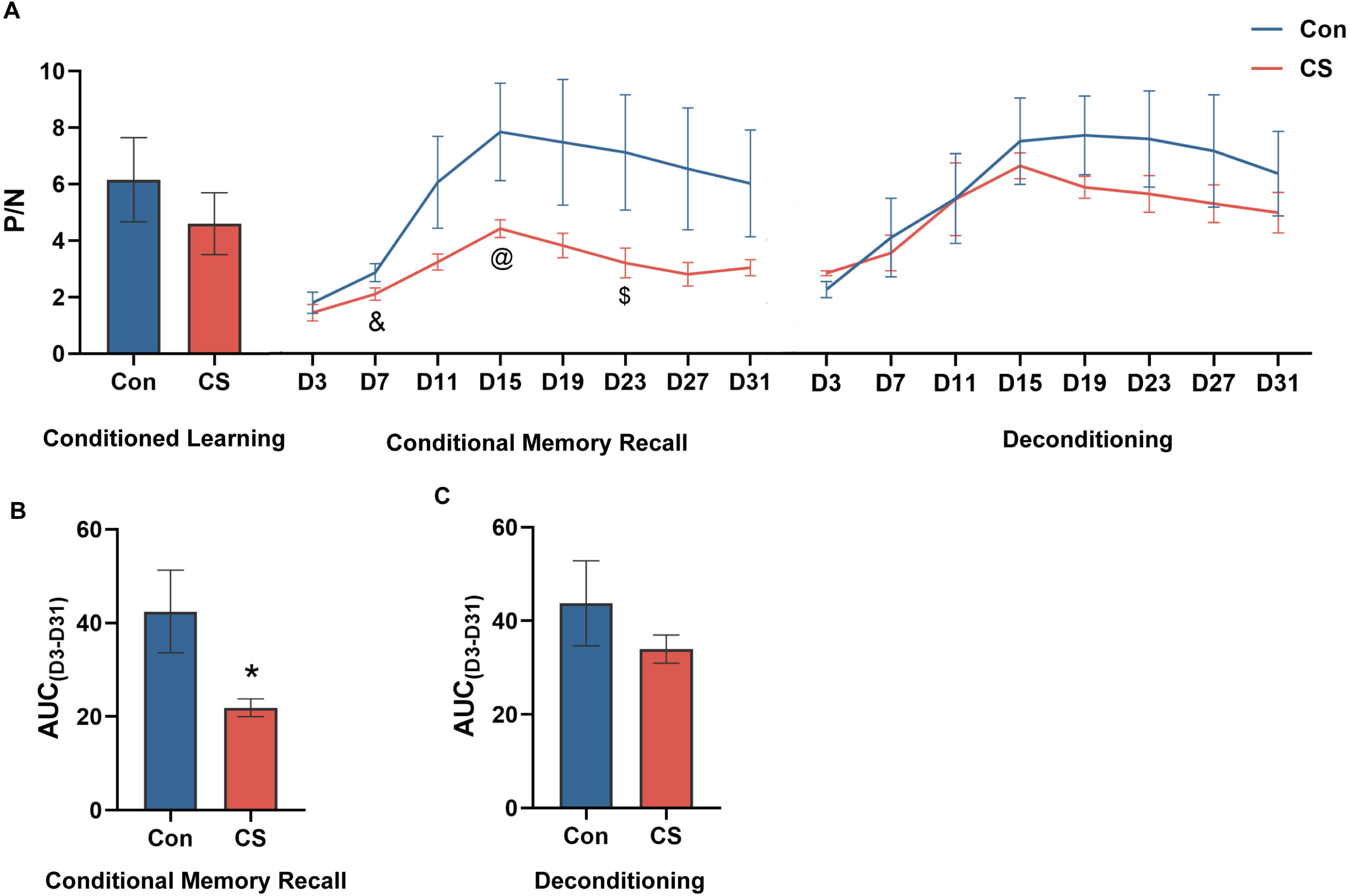
Associative learning of immune cells between PD and mcKLH-Rh1 conjugate. (A) The reactivity of immune serum to BSA-Rh1 conjugate (P/N) in the process of conditioned learning, conditional memory recall, and deconditioning. (B) Areas under curve (AUC) of P/N during conditional memory recall. (C) Areas under curve (AUC) of P/N during deconditioning. *p < 0.05 vs control group; & p = 0.085; @ p = 0.087; $ p = 0.054; mean ± SEM (n = 5).

## 4. Discussion

Associative learning is the ability of animals or human beings to perceive contingency relations between stimuli or events in their environment. It is a fundamental component of adaptive behavior as it allows anticipation of an event based on another. This adaptive behavior is usually used to assess cognitive function in animals or human beings[15, 16]. Besides vertebrates and invertebrates, a great deal of research on associative learning has been done even in plants and unicellular organisms (e.g., paramecia and amoeba)[17]. However, up to now, no studies on associative learning in cells in the body have been carried out. Initially, immortalized cell lines were adopted as research subjects to test the hypothesis proposed in this paper. We used low levels of endotoxin as the CS and hypoxia as the US, and measured cell survival to determine whether cell lines had established the CS-US association. However, cell health could not be maintained well throughout the processes of repeated hypoxia, passaging, and cryopreservation, and cell lines got susceptible to microbial infections, resulting in frequent interruptions of experiments.

Afterwards, we turned out attention to the immune cells in the body, since immunological memory is well known to be one of the cardinal features of the immune system (e.g., memory T cells, memory B cells, and memory-type natural killer cells)[18]. These memory immune cells are believed to persist long term in tissues and respond against a previously encountered antigen more quickly and better[19]. Thus, in the beginning, we planned to use yeast infection as the US, and had preliminarily observed the dynamic changes of defensive responses (e.g., body temperature, neutrophils, monocytes, and lymphocytes) in rats after repeated yeast infection. However, the research has been put on hold due to the lack of clarity on what conditioned stimulus to choose. Until recently, we successfully prepared anti-hapten polyclonal antibodies for the qualitative and quantitative detection of small-molecule compounds in brain tissues[11, 14], and finally conceived another experimental protocol to determine the presence of conditioned immune memory.

Based on hapten Rh1, we designed to use the analogues of Rh1 (e.g., PPD and PD) as the CS and the mcKLH-Rh1 conjugate as the US and determined the establishment of the CS-US association by detecting the dynamics of anti-Rh1 antibody production in mouse serum. In the study, PPD, which showed a cross-reaction with anti-Rh1 antibody by 29.7%, was preferred as the CS. As for the mice of conditioned stimulus group, intraperitoneal injection of PPD is repeatedly followed by subcutaneous injection of mcKLH-Rh1 conjugate. Whereas the control mice were repeatedly injected subcutaneously only with mcKLH-Rh1 conjugate. After the serum antibody response to Rh1 faded to a relatively stable level, the mice of two groups were intraperitoneally administered PPD at the same time. According to our assumptions, if the CS-US association was not established, two groups of mice would respond similarly against previously encountered mcKLH-Rh1 conjugate. Otherwise, the mice of conditioned stimulus group might respond against mcKLH-Rh1 conjugate more quickly or stronger than the control mice. However, the antibody response to Rh1 in the mice of conditioned stimulus group was unexpectedly slower and weaker than the control mice.

It was unlikely that the unexpected experimental phenomenon was due to the suppression of antibody response by PPD, as the control mice were also intraperitoneally administered PPD. In addition, immunosuppressive effects of PPD have not been reported in the literature[20, 21]. Thus, it could only be explained that immune cells recognized PPD, predicted the arrival of mcKLH-Rh1 conjugate, and made defensive behaviors of antibody response inhibition to alleviate the immune damage induced by mcKLH-Rh1 conjugate. Additionally, when the mice of two groups were directly injected with mcKLH-Rh1 conjugate without prior PPD administration, the difference between two groups in antibody response to the immunogen disappeared during deconditioning, suggesting that the immune cells have successfully established the CS-US association following multiple conditioned learning.

Next, we used PD as the CS that did not show any obvious cross-reaction with anti-Rh1 antibody. The results showed that the antibody response to Rh1 in the mice of two groups was like that in above studies regarding PPD as the CS throughout three phases of the experimental procedures. According to the recent literature, in addition to activities of anticancer, radio-resistance, and cholesterol homeostasis, PD did not exhibit immunosuppressive activity as well[20]. Consequently, this finding indicated that immune cells could successfully establish an association between Rh1 analogues and mcKLH-Rh1 conjugate, regardless of whether the analogue is like or different from the epitope of antigen. As far as the study is concerned, it is unknown which type of immune cells are involved in the process of conditioned immune memory, but associative learning itself suggests that the immune system in the body has a primary cognitive function including learning, memory, recognizing, and reasoning.

Before concluding, we should note that there are several limitations or shortcomings to the associative learning of cells in the body assessed in this study. First, the temporal coincidence between CS and US is crucial for associative learning. The inter-stimulus interval of the CS and US needs to fall within a brief coincidence time window[22]. In the study, the CS was given for three consecutive days immediately followed by the US, ensuring that the time interval between the CS and US is zero. Thus, the effect of coincidence time window on associative learning of immune cells in the body was not considered in the study. Second, we only tested serum antibody response to Rh1, not to mcKLH at the same time. Accordingly, during conditioned memory recall, it remains unclear whether the antibody response to mcKLH is consistent with the response to Rh1. Furthermore, as the levels of antibodies were not measured in the study, it is unknown whether the inhibited antibody response to Rh1 is due to decreased antibody production or a decrease in the affinity of antibodies to Rh1. Third, we observed that immune cells could associate Rh1 analogues (CS) with mcKLH-Rh1 (US) in the study. When anyone in all organic compounds is selected as the CS, whether immune cells can still establish the CS-US association is worthy of further investigation.

## 5. Conclusion

We demonstrated here that immune cells can acquire the new association between a hapten analogue and a hapten-protein conjugate after undergoing a classical conditioning procedure. The conditioned immunological memory enables immune cells to predict the arrival of an aversive immunogen and make a conditioned response that weakens humoral immunity to prevent immune damage. To our knowledge, this study is the first to discover the ability of immune cells to learn by associating. Our findings open a new frontier of exploring cellular cognitive functions and contribute to a novel understanding of bioactivities in the body. Additionally, the methodology of ‘abstraction and analogy’ in Chinese medicine can inspire us to explore and discover some immaterial aspects of life activity rules in the body.

## 6. Contributors

Zhu Zhu wrote the draft of the manuscript; Qin Wei and Qianlin Li undertook animal experiments; Hantao Zhang prepared hapten and performed ELISA examination; Zixuan Lin prepared the figures; Lin Chen assisted in the data analysis; Yunan Zhao conceived the idea, supervised, and supported the project; and all authors discussed the results and commented on the manuscript.

## Conflicts of interest

The authors have declared that there is no conflict of interest.

## Acknowledgments

The study was financially supported by National Natural Science Foundation of China (Nos. 82204647) and Jiangsu Provincial Administration of Traditional Chinese Medicine (Nos. MS2023006).

---

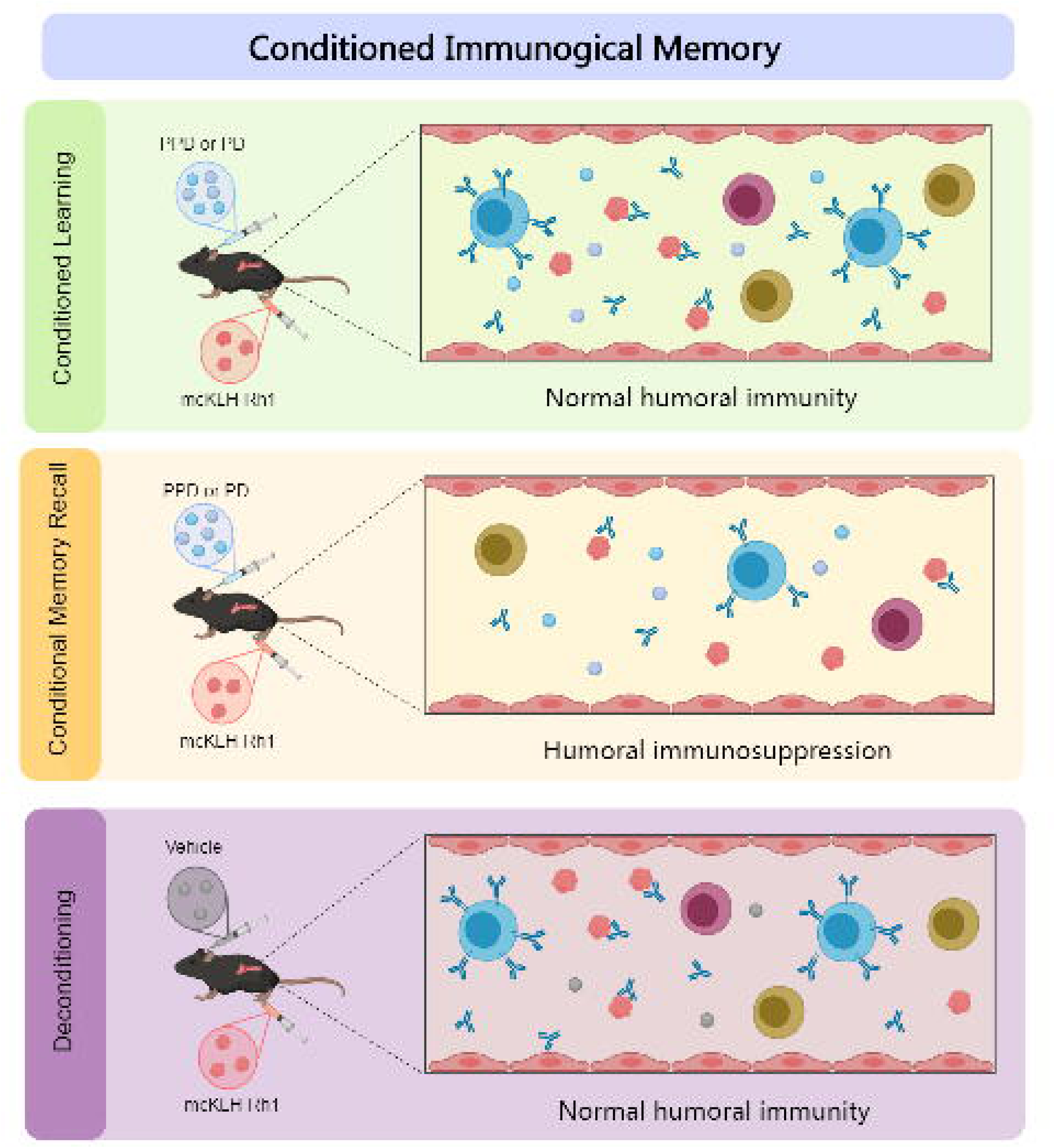

## Notes

### Competing Interest Statement

The authors have declared no competing interest.

